# Chilling, irradiation and transport of male *Glossina palpalis gambiensis* pupae: effect on the emergence, flight ability and survival

**DOI:** 10.1101/502567

**Authors:** Souleymane Diallo, Momar Talla Seck, Jean Baptiste Rayaissé, Assane Gueye Fall, Mireille Djimangali Bassene, Baba Sall, Antoine Sanon, Marc JB Vreysen, Peter Takac, Andrew G Parker, Geoffrey Gimonneau, Jérémy Bouyer

**Affiliations:** Centre International de Recherche-Développement sur l’Élevage en Zone Subhumide – CIRDES – Bobo – Dioulasso; Laboratoire d’Entomologie Fondamentale et Appliquée, Unité de Formation et de Recherche en Sciences de la vie et de la Terre, Université Ouaga I Pr Joseph Ki Zerbo, Burkina Faso; Institut Sénégalais de Recherches Agricoles, Laboratoire National d’Elevage et de Recherches Vétérinaires, Service de Bio-écologie et Pathologies Parasitaires, BP 2057, Dakar - Hann, Sénégal.; Direction des Services Vétérinaires – Dakar, Sénégal; Insect Pest Control Laboratory, Joint FAO/IAEA Programme of Nuclear Techniques in Food and Agriculture; Institute of Zoology, Section of Molecular and Applied Zoology, Slovak Academy of Sciences, Slovakia; Scientica, s.r.o., - Hybesova 33; Bratislava, Slovakia; CIRAD - Département Systèmes Biologiques - UMR 17 Intertryp CIRAD/IRD; CIRAD - Département Systèmes Biologiques - UMR ASTRE CIRAD/INRA

**Keywords:** Tsetse flies, mass-rearing conditions, sterile insect technique, quality

## Abstract

**Background:** The sterile insect technique (SIT) requires mass-rearing of the target species, irradiation to induce sexual sterility and transportation from the mass-rearing facility to the target site. Those treatments require several steps that may affect the biological quality of sterile males. This study has been carried out to evaluate the relative impact of the chilling, irradiation and transport on emergence rate, flight ability and survival of sterile male tsetse flies *Glossina palpalis gambiensis*.

**Results:** Chilling, irradiation and transport all affected the quality control parameters studied. The emergence rate was significantly reduced by long chilling periods and transport, i.e. from 92% at the source insectary to 78% upon arrival in Dakar. Flight ability was affected by all three parameters with 31% operational flies lossed between the source and arrival insectaries. Only survival under stress was not affected by any of the treatments.

**Conclusion:** The chilling period and transport were the main treatments which impacted significantly the quality of sterile male pupae. Therefore, the delivery of sterile males was divided over two shipments per week in order to reduce the chilling time and improve the quality of the sterile males. Quality of the male pupae may further be improved by reducing the transport time and vibration during transport.

**Author summary:** Tsetse fly and the disease it transmits, trypanosomosis, remain an enormous challenge in several countries in sub-Saharan Africa. The use of the Sterile Insect Technique (SIT) has become one of the components of the eradication of tsetse fly in Africa. The sterile insect technique (SIT) requires mass-rearing of the target species, irradiation to induce sexual sterility and transportation from the mass-rearing facility to the target site. In this study, we demonstrate the relative impact of the chilling, irradiation and transport on the emergence rate, the flight ability and the survival of sterile male tsetse flies *Glossina palpalis gambiensis*. We found that the chilling, irradiation and transport affected the emergence rate and flight ability. But the survival of the sterile male under stress was not affected by any of the treatments. Hence, the quality of the male pupae may further be improved by reducing the transport time and vibration during transport.

## Introduction

Tsetse flies (Diptera: Glossinidae) are the cyclical vectors of trypanosomes in sub-Saharan Africa, the causative agents of African animal trypanosomosis (AAT) or nagana for animals and human African trypanosomosis (HAT) or sleeping sickness in humans [1]. In the agricultural sector, the presence of tsetse flies limits the exploitation of fertile land in the more than 10 million km^2^ infested area and about 50 million cattle and tens of millions of small ruminants are permanently at risk of becoming infected by AAT [2]. This situation leads to high economic losses in the livestock sector estimated at USD 600–1200 million [3] and the overall annual losses in livestock and crop production are estimated at USD 4750 million [4]. Therefore, tsetse and trypanosomosis constitute a major constraint to livestock production and the main factor preventing the establishment of sustainable agricultural systems in much of sub-Saharan Africa. The lack of vaccines and high costs of disease treatment associated with the development of resistance by the parasites [5] make vector management a more reliable option for the control of the disease [6].

Currently, vector control can be achieved through several techniques, such as the sequential aerosol technique (SAT) and the ground spraying of insecticides, insecticide-treated targets or insecticide-treated animals as live baits, the use of traps and the sterile insect technique (SIT) [7], The principle of the SIT is to reduce the reproduction rate in a wild population by area-wide inundative releases of sterile male insects of the same species. The SIT has been used with success to suppress or eradicate several insect species of economic and sanitary importance [8]. For tsetse flies, the SIT has been used to eliminate populations of *Glossina palpalis gambiensis* Vanderplank *Glossina morsitans submorsitans* Newstead and *Glossina tachinoides* Westwood from 3000 km^2^ in Burkina Faso [9], and *Glossina palpalis palpalis* (Robineau-Desvoidy) from 1500 km^2^ in Nigeria [10]. Although these projects succeeded in eradicating the target populations, they were not conducted in the context of an area-wide integrated pest management (AW-IPM) approach and the cleared areas were thus re-invaded after the programs ended. The technology was, however, successfully applied within an AW-IPM approach to create a sustainable zone free of *Glossina austeni* Newstead (Diptera: Glossinidae) on Unguja Island of Zanzibar, United Republic of Tanzania in 1994-1997 [11]. This project confirmed the feasibility of integrating releases of sterile males with other suppression methods to create sustainable tsetse-free areas.

The Government of Senegal initiated a program in 2005 called “Projet d’éradication des mouches tsé-tsé dans les Niayes” [12] to eradicate *G. palpalis gambiensis* from a 1000 km^2^ area of the Niayes region neighbouring the capital, Dakar. The data generated by the feasibility study indicated the potential to create a sustainable zone free of *G. p. gambiensis* in the Niayes [12,13] and, therefore, the Government of Senegal opted for an AW-IPM approach including an SIT component. Since 2013, the sterile flies for the eradication campaign in Senegal have been provided by the Centre International de Recherche-Développement sur l’Elevage en zone Subhumide (CIRDES) in Bobo-Dioulasso, Burkina Faso, the Slovak Academy of Sciences (SAS) in Bratislava, Slovakia, and supplemented by the FAO/IAEA Insect Pest Control Laboratory (IPCL), Seibersdorf, Austria. In addition, since 2017, the Insectary of Bobo Dioulasso (IBD) in Burkina Faso has also provided sterile male pupae to the programme.

The supply of sterile flies required several important activities in the source insectary such as chilling and handling of the insect and their transport thereafter, that may have affected the quality of sterile males upon delivery at the Institut Sénégalais de Recherches Agricoles (ISRA) insectary, in Dakar.

Preliminary observations already showed that the flight ability of sterile male *G. p. gambiensis* sent to the ISRA insectary was low compared to flies emerged at CIRDES and SAS insectaries [14], and this was related to the chilling and irradiation treatments and the transport of the sterile pupae before they reached the ISRA insectary. This is of prime importance as the quality of the released sterile males remains one of the most crucial prerequisites for the success of AW-IPM programs that have an SIT component.

Based on the quality protocol described in Seck et al. [14], three biological parameters were measured to assess the impact of chilling, irradiation and transport on the quality of sterile males sent to Dakar for the eradication project in the Niayes area: i) adult emergence, ii) flight ability, and iii) survival of the flyers under starvation.

## Materials and Methods

### Insectaries

The study was carried out in three different insectaries between January 2015 and June 2016. Two were mass-rearing insectaries: the CIRDES insectary in Bobo-Dioulasso, Burkina Faso and the Slovak Academy of Sciences (SAS) insectary in Bratislava, Slovakia. The third one was the ISRA emergence insectary in Dakar, Senegal. The pupae and flies in these insectaries were kept under the same environmental conditions: 24–25°C, 75 ± 5% rH and 12:12 light/dark photoperiod.

### Biological material

Male pupae of the *G. p. gambiensis* BKF strain from the CIRDES and SAS colonies were used in this study. This tsetse colony has been maintained at the CIRDES insectary for more than 40 years and were offered blood meals using an *in vitro* silicon membrane feeding system using irradiated cow blood, collected from the local abattoir [15]. The colony was established in 1972 at MaisonAlfort (France) with pupae collected from Guinguette (Bobo-Dioulasso) and in 1975, the colony was transferred to the Centre de Recherche sur la Trypanosomiase Animale (CRTA) (later renamed CIRDES).

In 2009, 8000 pupae of this colony were shipped to the IPCL of the Joint FAO/IAEA Programme of Nuclear Techniques in Food and Agriculture to establish a colony for research purposes to support the eradication programme in the Niayes [16,17]. The IPCL colony provided seed material to the SAS where a colony was likewise established to supply additional pupae to the Senegal project.

### Chilling and irradiation

At the CIRDES and SAS insectaries, pupae were collected daily at the completion of female emergence and immediately chilled at 4°C to prevent male emergence and irradiated under chilled conditions (4–6°C) [18]. The CIRDES pupae were irradiated in a ^137^Cs source for 24 minutes and 30 seconds to yield a dose of 110 Gy. The SAS pupae were shipped chilled to the IPCL where they were irradiated in a ^60^Co irradiator (Gammacell 220, Nordion Ltd, Ottawa, Canada; dose rate of 131.4 Gy.min^-1^), or in a 150 kV X-ray irradiator (Rad Source RS2400; dose rate of 14.3 Gy.min^-1^).

### Packaging and transport of sterile pupae

After irradiation, pupae were packed and transported to Dakar as described in Pagabeleguem et al. and Seck et al. [14,18]. Briefly, irradiated pupae were placed in Petri dishes and packed in insulated boxes containing phase change material packs (S8, PCM Products Limited, Cambridgeshire, U. K.) to maintain the temperature between 10 ± 2°C. The box size and the number of S8 packs used were adjusted to the number of pupae shipped [18]. Pupae from CIRDES were then shipped with a courier service (DHL^®^) using public bus transport from Bobo-Dioulasso to Ouagadougou and commercial aircraft between Ouagadougou and Dakar. The average transport and chilling time for pupae from CIRDES was between 72 and 84 hours, that could be divided into 24 to 48 hours chilled at 8 ± 2°C in the source insectary and ±36 hours at 10 ± 2°C during the transport to Dakar.

### Effect of chilling, irradiation and transport on male pupae performance parameters

The objective of the quality control is to evaluate the quality of sterile males, measured by the percentage of operational flies corresponding to the percentage of flies escaping the flight device [14]. Here, these quality control parameters were used to measure the impact of various treatments (chilling, irradiation, transport) on the emergence rate and rate of operational flies. According to the successive steps set up to prepare and ship pupae, five treatments were defined for the CIRDES pupae:

- A0 (CIRDES_A0) = 1 sample of 50 pupae (the control group): no chilling and no irradiation (equivalent to the conditions of the colony);

- A1 (CIRDES_A1) = 1 sample of 50 pupae: chilling (8 ± 2°C) of the A0 pupae for 24 to 72 hours (no irradiation);

- A2 (CIRDES_A2) = 1 sample of 50 pupae: irradiation of A1 pupae;

- A3 (CIRDES_A3) = 1 sample of 50 pupae: second chilling (8 ± 2°C) of A2 pupae for 48 hours.

A temperature and humidity Hobo® data logger (Hobo® model EL-USB-2) was placed in the chiller to record the temperature and relative humidity;

-A4 (CIRDES_A4) = a final group of 50 pupae was tested at ISRA after transport. These pupae accumulated all the treatments of A3 and the transport (by road and air) in an insulated box to maintain the temperature between 10 ± 2°C. A temperature and humidity Hobo® data logger (Hobo® model EL-USB-2) was added to the shipping box during the packing at CIRDES to record the temperature and relative humidity inside the insulated box every 5 minutes.

For the SAS pupae, the quality control tests were only performed for the A1 treatment (SAS_A1).

For each treatment, a standard quality control protocol was applied. Briefly, the 50 pupae were put in Petri dishes under ~1cm of sand mixed with a fluorescent dye (DayGlo) (0.5g dye/200g of sand), to mimic the natural emergence conditions in the soil and to allow discrimination of the sterile male flies from wild flies during an entomological monitoring in AW-IPM programmes that have an SIT <u>component</u>. The Petri dish was then put in a flight cylinder, i.e. PVC tube 10 cm high and 8.4 cm in diameter [14]. The inner wall of the cylinder was coated with unscented talcum powder to prevent the flies from crawling out. Flies flying out of the tube were considered as “operational flies” (i.e. available for SIT).

The survival of the sterile males that escaped the flight cylinder was assessed under stress conditions (no blood meal). Every morning, the emerged flies were collected and transferred to standard fly holding cages. The flies emerged on a given day were pooled in one cage. Dead flies were counted daily and removed from the cages.

This quality control protocol was implemented between January 2015 and June 2016 with samples of pupae collected from each batch sent to Dakar for the eradication program (twice per week from CIRDES and once per week from SAS).

### Data analysis

Data analysis was performed using the R software version 3.5.1 [19]. Data were analysed using binomial linear mixed effects models using the package lme4 [20], with the emergence rate or rate of flyers as the response variables, the batch origin (CIRDES or SAS) and treatment type (A0 to A4) as fixed effects and the date of arrival at ISRA as a random effect [21,22]. A Gaussian linear mixed effect model was used to analyse the mean survival under starvation, with the same fixed and random effects as in the previous models.

## Results

### Emergence rate

The mean emergence rate observed for the five treatments ranged between 92% for A0 and 78% for A4 with an overall mean at 83% (Table 1, Figure 1). The model results showed that at the CIRDES, the first and second chilling rounds (A1 and A3) had a significant negative effect on the emergence rate (*P* < 0.001) whereas irradiation (A2) did not significantly reduce it further (*P* > 0.05; Table 2). The rate of emergence was also superior for A1 at CIRDES than SAS (*P* < 0.001), although the mean values were very close (Table 1). The transport also significantly reduced the rate of emergence (*P* < 0.001; the reference level was set to CIRDES_A3 to calculate this probability).

**Figure 1.**
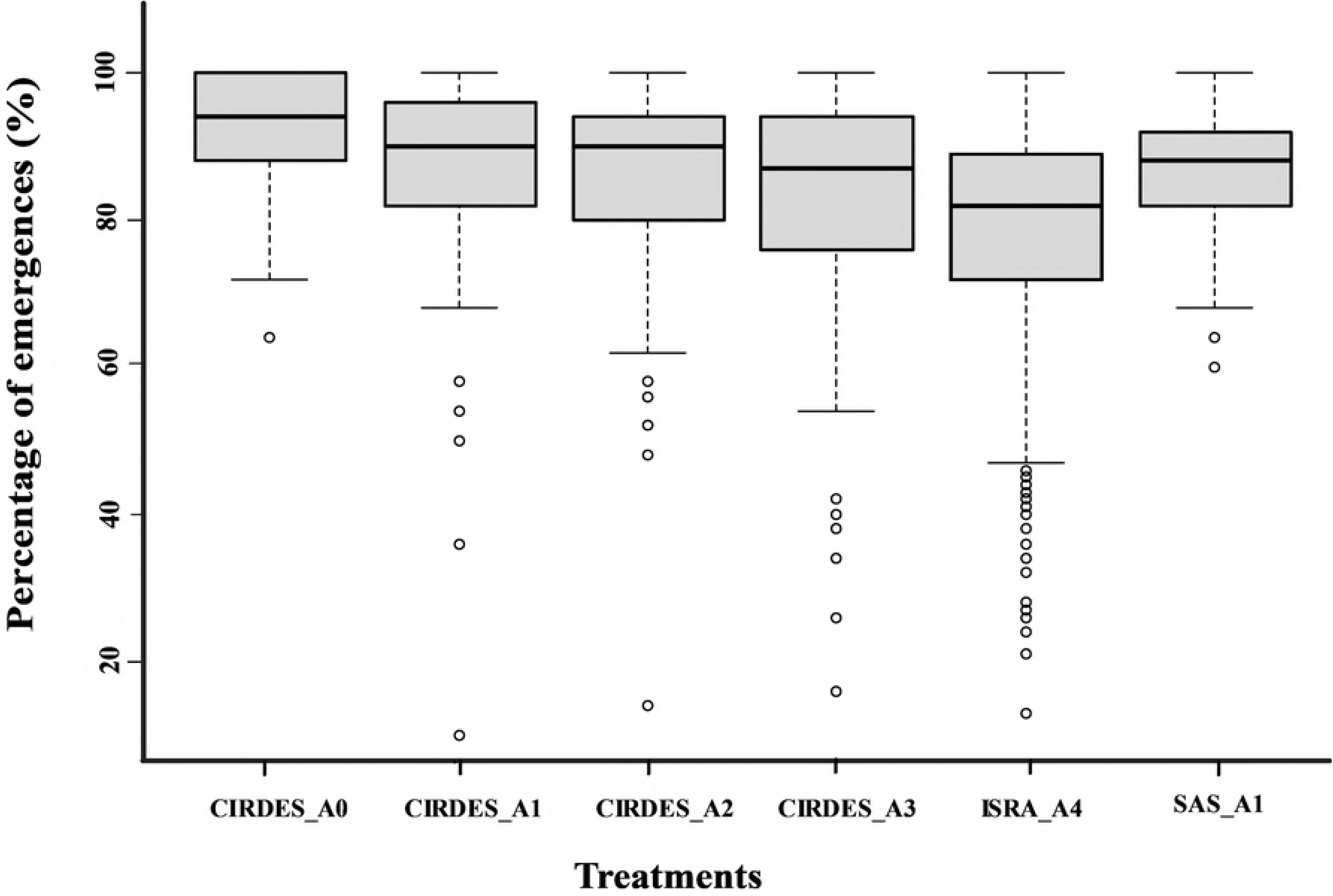
Percentage of emergence according to the treatment and sites. Boxes extend between the 25^th^ and 75^th^ percentile. A thick line denotes the median. The whiskers extend up to the most extreme values and white circle represents outliers data.

**Table 1.** Average percentages of emergence and operational flies depending on the treatment (A0 to A4) and the site where the test was performed. Batches correspond to the number of subsamples of 50 pupae collected from each consignment sent to Dakar.

| Site test | Treatment | Nb batches | Mean emergence rate (±SD) | Mean operational rate (±SD) |
| --- | --- | --- | --- | --- |
| CIRDES | A0 | 116 | 92±8 | 82±13 |
| CIRDES | A1 | 110 | 87±13 | 64±20 |
| CIRDES | A2 | 113 | 85±13 | 45±21 |
| CIRDES | A3 | 114 | 82±17 | 52±21 |
| ISRA | A4 | 468 | 78±15 | 51±21 |
| SAS | A1 | 146 | 86±8 | 82±9 |
| TOTAL | | 1067 | 83±14 | 60±23 |

NB= number; SD= Standard Deviation

**Table 2.** Summary of the binomial linear mixed effects models for emergence rate. The

| Fixed effects | Estimate | Std. Error | Z value | P value |
| --- | --- | --- | --- | --- |
| Intercept | 2.249 | 0.063 | 35.76 | <0.001 |
| CIRDES A0 | 0.585 | 0.065 | 9.06 | <0.001 |
| CIRDES A2 | -0.104 | 0.056 | -1.86 | 0.0635 |
| CIRDES A3 | -0.393 | 0.054 | -7.24 | <0.001 |
| ISRA A4 | -0.887 | 0.045 | -19.51 | <0.001 |
| SAS A1 | -0.43 | 0.059 | -7.24 | <0.001 |

### Operational flies

The highest mean rate of operational flies was observed for treatments A0, and A1 SAS with a value of 82% and the lowest was observed for the treatment A4 (pupae upon arrival at the Dakar insectary) with 51% (Table 1 and Figure 2). Results of the binomial mixed effects model showed that the first chilling round (A1) and the irradiation (A2) both reduced the quality of the flies at CIRDES (*P* < 0.001; Table 3). For A1 pupae, the rate of operational flies was significantly better for the SAS flies than the CIRDES flies (*P* < 0.001). Interestingly, the second chilling event (A3) induced a significant “recovery” of the rate of operational flies at the CIRDES (*P* < 0.001, the reference level was set to CIRDES_A2 to calculate this probability). Finally, the transport from the CIRDES to Dakar also significantly reduced the quality of the sterile males (*P* < 0.001, the reference level was set to CIRDES_A3 to calculate this probability) compared with sterile males emerged from the A3 treatment (Figure 2).

**Figure 2.**
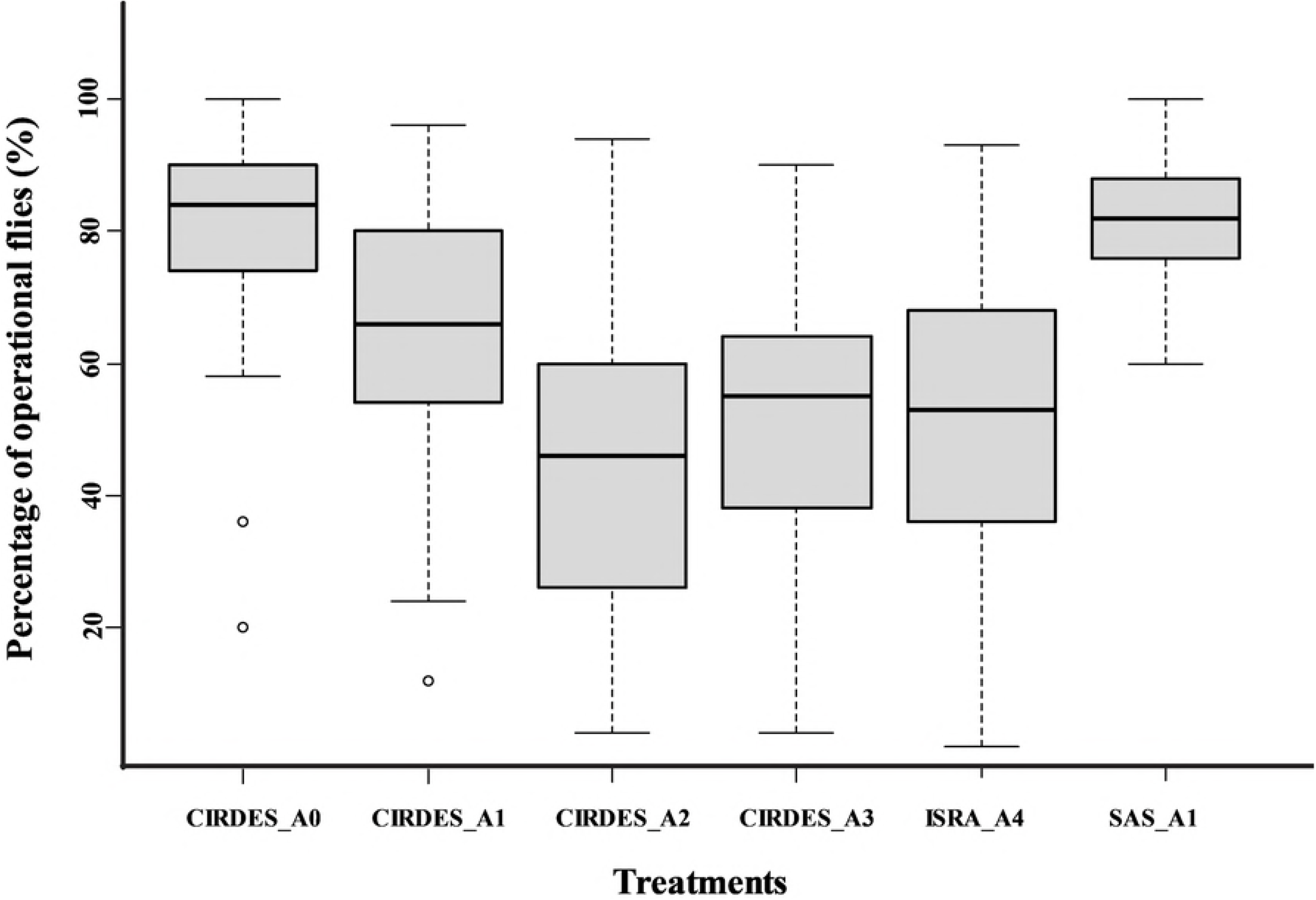
Percentages of operational flies (%) according to treatment (A0 to A4) and site where the test was performed. Boxes extend between the 25^th^ and 75^th^ percentile. A thick line denotes the median. The whiskers extend up to the most extreme values, and white circles represents outliers data.

**Table 3.** Summary of the binomial linear mixed effects models for the operational rate. The reference level is CIRDES_A1.

| Fixed effects | Estimate | Std. Error | Z value | P value |
| --- | --- | --- | --- | --- |
| Intercept | 0.784 | 0.053 | 14.74 | <0.001 |
| CIRDES A0 | 1.001 | 0.046 | 21.77 | <0.001 |
| CIRDES A2 | -0.813 | 0.04 | -20.24 | <0.001 |
| CIRDES A3 | -0.542 | 0.04 | -13.55 | <0.001 |
| ISRA A4 | -0.737 | 0.033 | -21.95 | <0.001 |
| SAS A1 | 0.673 | 0.047 | 14.17 | <0.001 |

### Survival under stress

Fly survival rates under starvation were very similar between treatments ranging from 4 to 5 days, except for A1 SAS that showed a mean survival rate of 2.34 days (Figure 3). Model results showed that at the CIRDES none of the treatments had an impact on survival (Table 4; *P* > 0.05; the reference level is CIRDES_A1). However, the mortality rate was much higher for the SAS flies as compared with the CIRDES flies for the A1 treatment group (*P* < 0.005). Finally, the survival of the A4 batch at the ISRA (after transport) was significantly better than the A3 batch at the CIRDES (*P* = 0.006, the reference level was set to CIRDES_A3 in the previous model to calculate this probability).

**Figure 3.**
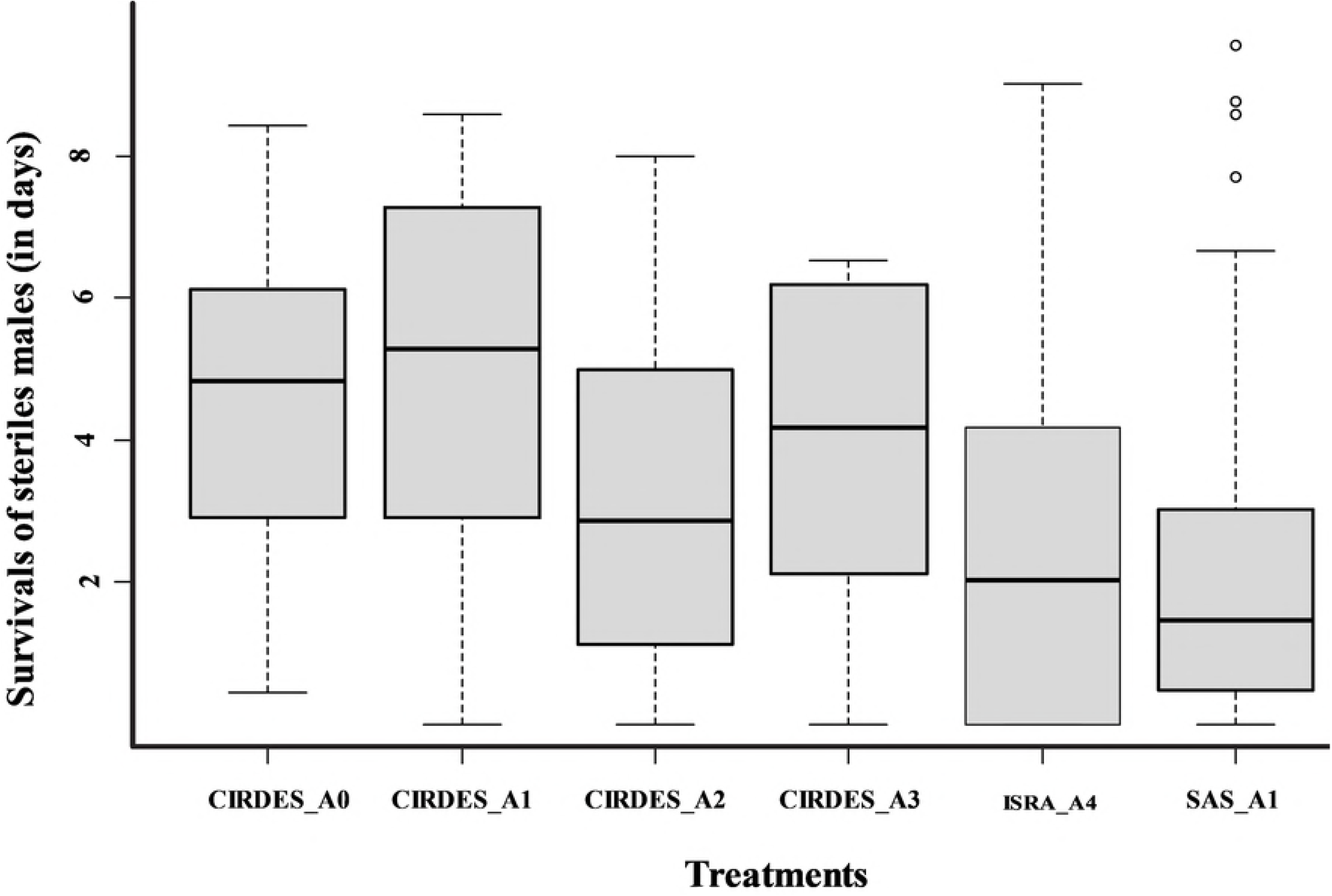
Boxplots of the survival of sterile males (in days) monitored under starvation conditions during the quality test for the four treatments (A0 to A4) and the three insectaries where the tests <u>were carried out</u>. Boxes extend between the 25^th^ and 75^th^ percentile. A thick line denotes the median. The whiskers extend up to the most extreme values, and the white circles represent outlier data.

**Table 4.** Summary of the linear models for mortality rate. The reference level is CIRDES_A1

| Fixed effects | Estimate | Std. Error | Z value | P value |
| --- | --- | --- | --- | --- |
| Intercept | 4.35 | 0.104 | 41.687 | <0.001 |
| CIRDES A0 | 0.244 | 0.14 | 1.742 | 0.082 |
| CIRDES A2 | 0.023 | 0.149 | 0.154 | 0.8779 |
| CIRDES A3 | 0.183 | 0.147 | 1.24 | 0.2151 |
| ISRA A4 | 0.491 | 0.111 | 4.414 | <0.001 |
| SAS A1 | -2.01 | 0.153 | -13.132 | <0.001 |

## Discussion

The sterile insect technique depends on the mass-production of sterile male insects of good biological quality to be released into the target wild populations. The tsetse eradication programme in the Niayes of Senegal is implemented following an AW-IPM approach with a SIT component and aims at eradicating the native *G. p. gambiensis* populations in that region [23]. The sterile insects are provided to the project as irradiated pupae that are being transported from three production centres (two in Burkina Faso and one in Slovakia, Europe) under chilled conditions. Handling, irradiation, chilling and transport may affect the quality of the adult male flies and their performance upon arrival in Dakar. Previous studies have already shown that the emergence rate and the quality of the sterile males are lower in the Dakar insectary than in the production centres [14,18].

### Emergence rate

The emergence rate of pupae was negatively affected by all handling steps, decreasing from 92% for A0 pupae at the CIRDES insectary to 78% for A4 pupae at the ISRA insectary. Both chilling steps (A1 and A3) significantly reduced the number of emerging flies whereas irradiation (A2) did not. This result contrast with previous studies on the chilling effect on tsetse fly pupae. Mutika et al. [16] worked with the same *G. p. gambiensis* BKF strain and observed that emergence of 28-day old male pupae after storage at low temperature (10°C) for 3, 5, or 7 days was similar to those kept under standard colony conditions. Similar results were observed for pupae of *G. morsitans* maintained at 12°C for two weeks [24]. Our results of the current study indicate that chilling affected tsetse emergence rate significantly and the chilling period must be kept as short as possible. It must be noted that our results were obtained with a temperature that was on average 2 to 4°C lower than the previous studies, which might partially explain this discrepancy. On the other hand, irradiation had no significant negative effect on pupal emergence rate. Similar results were observed for *G. tachinoides* pupae irradiated with 120 Gy on day 28 post larviposition [25] and for *G. p. gambiensis* pupae irradiated with 110 Gy [16]. These results were expected since several studies in the past have aimed at optimising the irradiation process in order to induce near 100% sterility while minimising somatic damage [26,27].

The last step, transport to Dakar, also significantly reduced fly emergence rate. During shipment, pupae are exposed to uncontrolled factors such as vibrations or mechanical shocks that are absorbed by pupae and may affect the fly emergence rate. Therefore, solutions could be tested such as the use of cotton wool, sawdust or vermiculite to cushion the mechanical shocks or vibrations during transport. Moreover, adult emergence may also be affected by excessive temperatures or inappropriate relative humidity during the rearing process [28]. Although adult emergence was negatively affected by the various treatments, the emergence rate improved with time during project implementation. Indeed, Pagabeleguem et al. [18] observed an emergence rate of pupae received at the ISRA of 74 ± 13.9% for shipments between January 2011 and 2014 against 78 ± 15% in this study. Moreover, our result was not different from those of Mutika et al. [16], who observed emergence rates between 76 to 91% for irradiated pupae chilled for 5 days.

### Operational flies

The rate of operational flies was also affected by all pupal treatments, decreasing from 82 ± 13% for A0 pupae to 51 ± 0.21% for A4 pupae. For the two first steps A1 and A2, 18% and 19% of operational flies are lost respectively, leading to a cumulative loss of 47% after these two steps.

These results contrast with several studies that showed that storage of mature pupae at low temperature (up to 5 days between 7-10°C) has no detrimental effects on male emergence, flight ability or survival [16,24,29,30]. The chilling period for A1 and A2 pupae were comparatively short (only 24 and 72 hours at 8°C) and such losses could not be explained by the cold exposure only. Reducing the chilling period could be one lever to improve these results.

Moreover, the rate of operational flies was significantly better for SAS pupae compared to CIRDES pupae of treatment A1. This difference could be explained by the different chilling durations between the SAS and the CIRDES pupae, given that CIRDES pupae were chilled for two to three days before their irradiation whereas it is only one to two days for the SAS pupae. Therefore, chilling duration and temperature seems to be the two most important parameters that must be surveyed continuously in order to improve fly quality. Although irradiation (A2) also affected the rate of operational flies, it may be due to the irradiation handling process instead of the radiation dose itself. Earlier studies have examined the effect of radiation dose on quality of pupae and they found that there is no effect on biological quality of pupae when the irradiation is done close to emergence [16,26].

Interestingly, the second chilling event (A3) led to a significant “recovery” of the rate of operational flies at the CIRDES compared to the A2 treatment and this effect was also confirmed upon arrival in Dakar (A4). This might be due to a positive impact of the reduced metabolic rate due to the chilling on cellular mechanisms to repair somatic damage.

The long shipment to Dakar also significantly affected the rate of operational flies, as already demonstrated [14,18]. Physical injuries during transport are probably responsible for this loss. Those physical injuries could the vibrations during the transport. We suggest a new study on the effect of vibration on tsetse flies irradiated pupae during the transport. However, as for the emergence rate, the results observed here seem to improve during project implementation. Indeed, Seck et al. [18] observed a rate of operational flies at the ISRA of 35.8± 18.4% for shipments between May 2012 to January 2015 against 51 ± 21% in this study. This increase shows that the overall handling process of flies since its first implementation was continuously improved and could probably be improved even more.

### Survival under stress

None of the treatments had an impact on fly survival (*P* > 0.05). However, the mean survival was significantly lower for the SAS pupae as compared with the CIRDES pupae for the A1 treatment group (P < 0.001), which is probably due to different quality of the blood diet and female performance in the two insectaries. Indeed, during this experiment, newly emerged sterile males were not offered a blood meal, and their survival only depended on the fat reserves acquired during larval development. This fat reserve is closely related to the quality and quantity of blood meals that are taken by the female parent and also to factors which affect the amount of fat consumed during pupal development and hence influence the lifespan of the newly emerged fly [31].

The survival of the A4 batch at the ISRA (after transport) was significantly better than the A3 batch at the CIRDES (*P* = 0.006) which here again suggest that the environmental conditions at the ISRA insectary were more suitable for *G. p. gambiensis* than at the CIRDES. In this case, blood could not be involved because sterile male flies originated from the same insectary.

## Conclusion

This study highlights the effects of several steps necessary to prepare and ship sterile male tsetse pupae in AW-IPM programs that have an SIT component. Chilling, irradiation and transport all negatively impacted the quality of sterile male pupae with lower emergence and operational rate of flies, but no effect was observed on survival. Although all these processes negatively impact the sterile male yield, longitudinal comparison of data from the Niaye erradication project highlight that sterile male quality improved with time. In order to limit the chilling effect, the project decided to split the delivery of sterile males into two shipments per week in order to reduce the chilling time and improve the quality of sterile males. Further optimizing the quality of the sterile male tsetse could be obtained by reducing the effects of chilling and transport vibration. It would be of interest to determine the threshold of chilling duration at which the emergence rate and the rate of operationnal flies can be optimised.

## Acknowledgement

We thank technicians of CIRDES, ISRA and SAS insectaries for their technical support.

Authors email addresses
Souleymane Diallo, Momar Talla Seck, Jean Baptiste Rayaissé, Assane Gueye Fall , Mireille Djimangali Bassene , Baba Sall, Antoine Sanon, Marc JB Vreysen, Peter Takac, Andrew Gordon Parker, Geoffrey Gimonneau , Jérémy Bouyer

